# Nick-seq for single-nucleotide resolution genomic maps of DNA modifications and damage

**DOI:** 10.1101/845768

**Authors:** Bo Cao, Xiaolin Wu, Jieliang Zhou, Hang Wu, Michael S. DeMott, Chen Gu, Lianrong Wang, Delin You, Peter C. Dedon

## Abstract

Here we present the Nick-seq platform for quantitative mapping of DNA modifications and damage at single-nucleotide resolution across genomes. Pre-existing breaks are blocked and DNA structures converted to strand-breaks for 3’-extension by nick-translation to produce nuclease-resistant oligonucleotides, and 3’-capture by terminal transferase tailing. Libraries from both products are subjected to next-generation sequencing. Nick-seq is a generally applicable method illustrated with quantitative profiling of single-strand-breaks, phosphorothioate modifications, and DNA oxidation.

## Main text

Genomic mapping of specific DNA modifications^1^ and damage^2^ can be achieved with methods such as bisulfite sequencing for 5-methylcytidine^3^, chromatin immunoprecipitation (ChIP) coupled with next-generation sequencing (NGS)^4,5^, and single-molecule real-time (SMRT)^6^ and nanopore^7^ sequencing. However, all are limited to specific modifications or suffer from low sensitivity and specificity. Here we describe Nick-seq for highly sensitive quantitative genomic mapping of any type of DNA modification or damage that can be converted to a strand-break. As shown in Figure 1a, purified genomic DNA is subjected to sequencing-compatible fragmentation and the resulting 3’-OH ends are blocked with dideoxyNTPs. The DNA modification is then converted to a strand-break by enzyme or chemical treatment, followed by capture of the 3’- and 5’-ends of resulting strand-breaks using two complementary strategies. One portion of DNA is subjected to nick translation (NT) with α-thio-dNTPs to generate 100-200 nt phosphorothioate-containing oligonucleotides that are resistant to subsequent hydrolysis of the bulk of the genomic DNA by exonuclease III and RecJ_f_. The purified PT-protected fragment is used to generate an NGS library with the modification of interest positioned at the 5’-end of the PT-labeled fragment. A second portion of the same DNA sample is used for terminal transferase (TdT)-dependent poly(dT) tailing of the 3’-end of the strand-break, with the tail used to create a sequencing library by reverse transcriptase template switching^8^. Subsequent NGS positions the modification of interest 5’-end of the poly(dT) tail.

**Figure. 1.**
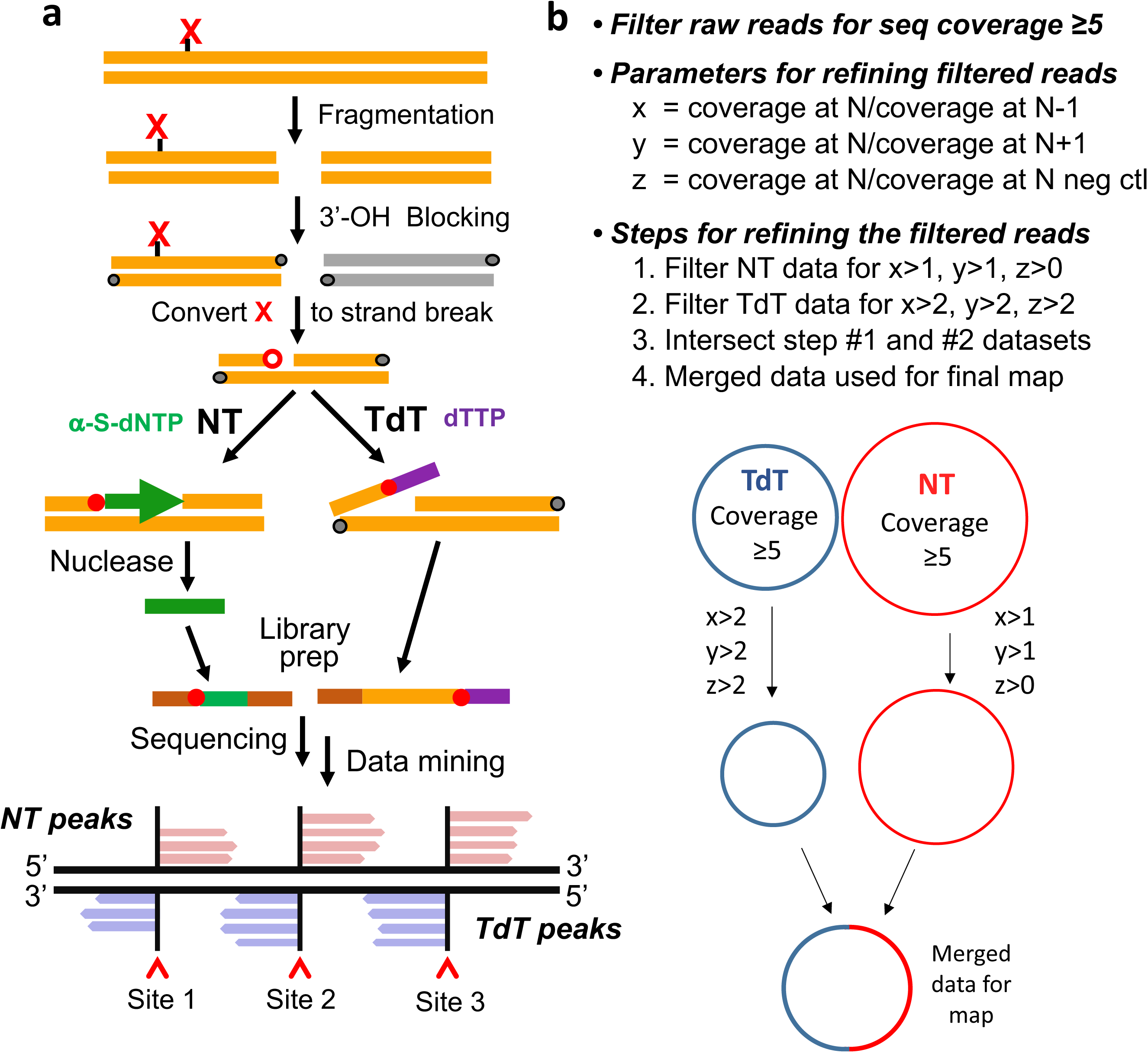
Nick-seq library preparation (a) and data analysis workflow (b).

The workflow for sequencing data processing (Fig. 1b) uses the NT-derived reads as the primary dataset for developing a rough modification map, with TdT-derived reads as complementary corrective data. This hybrid approach exploits the fact that NT is agnostic to the base identify at the damage site but generates a high background of false positive sites, while TdT cannot be used with modifications occurring at dT due to loss of the poly(dT) tail during data analysis. The TdT reads are used to correct NT false-positive reads. For example, if NT maps a strand-break at T 1000 in the genome, then TdT reads are examined for a strand-break at position 999 and 1001. This 1 position shift accommodates poly(dT) tail removal during data processing and validates the NT map. If TdT does not call a strand-break at 999 or 1001, then the NT result is considered a false positive. In other cases, if the NT-detected site occurs at a G, C, or A, then TdT valdiates the same site. The use of both methods increases the sensitivity and specificity of the resulting map.

We validated Nick-seq by mapping DNA single-strand-breaks caused by the site-specific endonuclease, Nb.BsmI, at the 2,681 G/CATTC motifs in the *E.coli* genome. Purified DNA was treated with Nb.BsmI and the Nick-seq-processed library sequenced using the Illumina NextSeq platform with an average of 10^7^ raw sequencing reads for each sample (Fig. 2b). Paired-end sequencing confirmed that >80% of reads uniquely aligned to the *E.coli* genome (**Supplementary Table 1**). For subequent reads enrichment (Fig. 1b), we calculated position-wise coverage values using the 5’-end of sequencing reads (NT read 1, TdT read 2) and defined Nick-seq peaks as having >5 reads and 2-times more reads than sites located one-nucleotide up- and down-stream. We then calculated the coverage ratio of the peaks to corresponding sites in an untreated DNA control. To identify the optimal minimal coverage ratio, we varied the ratio and calculated the number of identified sites at each ratio value (Fig. 2b). As the coverage ratio increased from 2 to 7, the number of identified sites decreased from 92% to 59% of 2,681 expected (“sensitivity”), while the accuracy (identified sites/expected sites; “specificity”) only increased from 98% to 99.5%. To maximize sensitivity, we chose a coverage ratio of 2, which allowed identification of 2,462 (97.5%) of the predicted Nb.BsmI sites. Another 1% of called sites (27) occurred in sequences differing from the consensus by one nucleotide (**Supplementary Table 2**). These sites showed lower average sequencing coverage (75 vs 1318) and likely represent Nb.BsmI “star” activity. Another validation experiment with Nb.BsrDI, which cuts at N/CATTGC, showed no evident 3’-end sequence bias for DNA break site detection (**Supplementary Table 3)**. Thus, Nick-seq showed high accuracy and sensitivity for single-nucleotide genomic mapping of DNA strand-breaks.

**Figure. 2.**
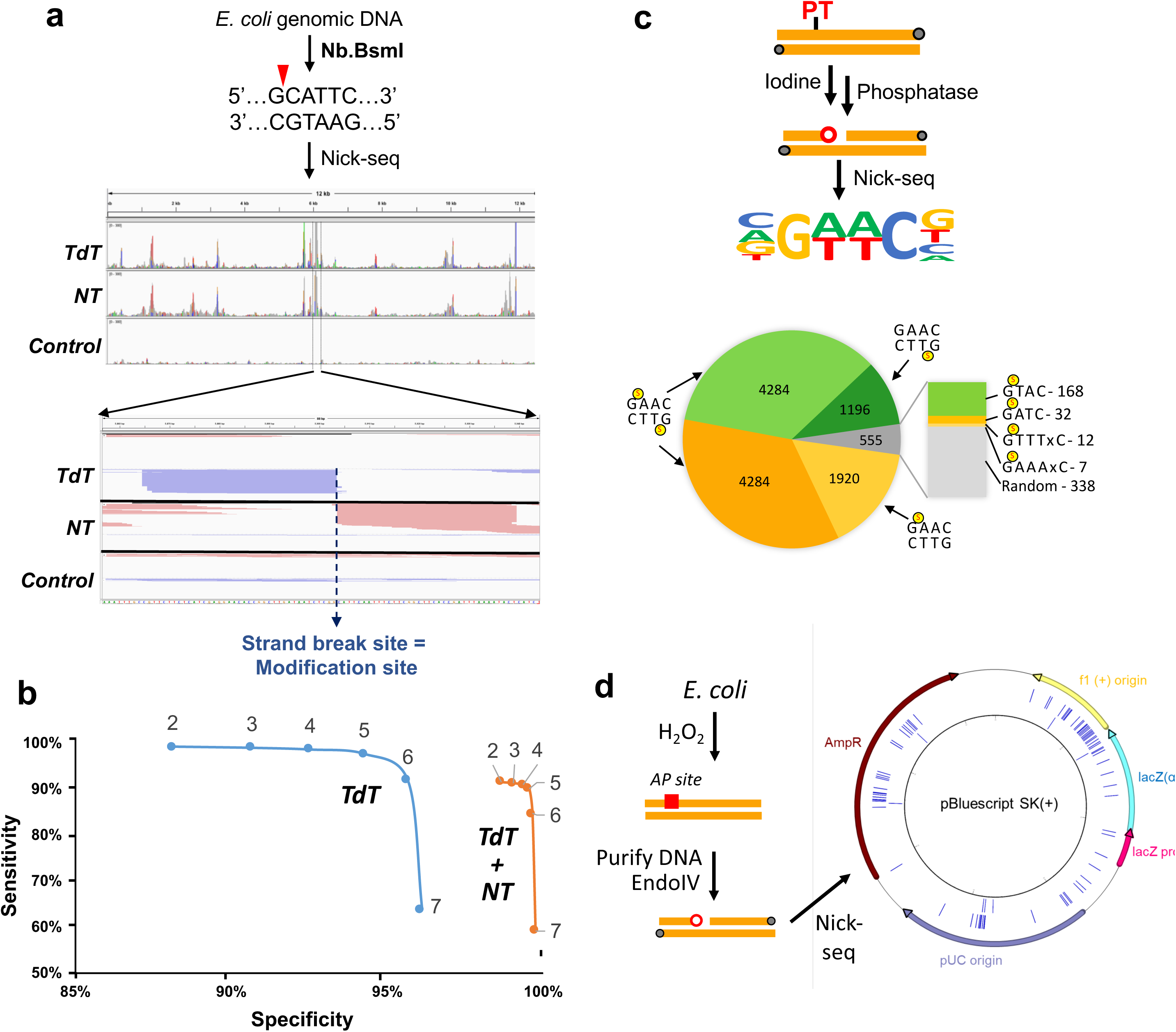
Nick-seq validation and application. (**a**) Mapping single-strand breaks produced by Nb.BsmI in *E. coli* genomic DNA. Middle panel: representative view of sequencing reads distirbuted in one genomic region. Red and green peaks mark reads mapped to forward and reverse strands of the genome, respectively. Lower panel: amplification of the region surrounding one peak, with read pile ups for TdT and NT sequencing converging on the site of the strand-break. (**b**) Nb.BsmI mapping data were used to define data mining parameters for accuracy and sensitivity of Nick-seq. In general, higher ratios yield greater accuracy but lower sensitivity. (**c**) Mapping PTs across the *S. enterica* genome by Nick-seq. (**d**) Application of Nick-seq to quantify abasic sites generated by H_2_O_2_ exposure in *E. coli*.

The validated Nick-seq was applied to map the naturally-occurring phosphorothioate (PT) DNA modifications in *Salmonella enterica* serovar Cerro 87, using iodine to oxidize PTs to produce DNA strand-breaks^9^ (Fig. 2c). We previously established by SMRT sequencing that PTs occurred as bistranded modifications at 10-15% of the GAAC/GTTC motifs in *S. enterica*^9^. Nick-seq recognized 12,239 PT sites (Fig. 2c, **Supplementary Table 4**), of which 11,684 (96%) occurred at G_PS_AAC/G_PS_TTC, with 8,568 (73%) modified on both strands and 27% modified on one strand (Fig. 2c). This agrees with our previous observations using an orthogonal sequencing method^9^. In addition to GAAC/GTTC motifs, Nick-seq also revealed less abundant PTs at G_PS_TAC (168), GPSATC (30), and G_PS_AAAC or G_PS_AAAAC (19), with half of G_PS_TAC and G_PS_ATC sites modified on only one strand (**Supplementary Table 4**). These results indicate that Nick-seq has a higher sensitivity to detect rare PT modifications than other methods.^9, 10^

Finally, we applied Nick-seq to DNA modifications not previously subjected to genomic mapping: oxiatively-induced abasic sites. Apurinic and apyrimidinic (AP) sites represent a prevalent and toxic form of DNA damage that blocks DNA replication and transcription^11, 12^. AP sites arise as intermediates in base excision DNA repair, in which damaged bases are excised by glycosylases and the resulting AP sites cleaved by AP endonucleases^13^. AP sites can also arise by oxidation of DNA on both the nucleobase and 2’-deoxyribose moietities^14, 15^, as well as by demethylation of m^5^C epigenetic marks.^16^ In spite of the importance of AP sites, little is known about their formation, persistence, and distribution in genomic DNA. Here we used Nick-seq to profile AP sites in *E. coli* exposed to H_2_O_2_ at a non-lethal dose of 0.2 mM (LD_50_ ~5 mM) (Fig. 2d). Following DNA purification, AP sites were expressed as strand-breaks using endonuclease IV (EndoIV), which cleaves both native and oxidized AP sites^14, 15^. Nick-seq identified 1,519 EndoIV-sensitive sites, as well as 82 sites in the endogenous plasmid, with an unexposed control showing 11 and 8 sites, respectively (Fig. 2d, **Supplementary Tables 5, 6**). Considering the nucleobase precursor of the AP site, there was a weak preference for thymine (33%) followed by adenine (25%), cytosine (24%), and guanine (18%), with a similar distribution in the plasmid (**Supplementary Tables 5, 6**). This suggests that H_2_O_2_-derived DNA oxidizing agents either do not selectively oxidize guanine as predicted^17^ or that the predominant form of damage is DNA sugar oxidation. However, there was a more pronounced sequence context effect. Analysis of 15 bp up- and down-stream of the AP sites revealed a strong preference for cytosine (47%) at −1 relative to the AP sites. The distribution of AP sites on the plasmid was also non-random (Fig. 2d), with clustering in three regions related to DNA replication and transcription: the F1 origin, pUC origin, and Amp^R^ gene (**Supplementary Table 7**). AP site clustering near DNA replication sites was observed previously by immunostaining^18^, suggesting that the transcriptionally active and single-strand DNA are vulnerable to oxidatively-induced AP sites. We tested this by analyzing the distribution of AP sites in the *E. coli* genome relative to origins of replication (OriC), coding sequences, and non-coding sequences. While there was an average of 0.32 Aps/kbp (1,519 AP per 4,686,000 bp), the 20 kb region around OriC showed 0.70 Aps/kbp. Nick-seq also revealed 1401 APs in the 4.1×10^6^ bp coding sequence region (0.34 AP/kbp), and 118 APs (0.20 AP/kbp) in the 0.58 x10^6^ bp non-coding region. These results suggest a preference for AP sites in DNA undergoing replication or transcription during H_2_O_2_ stress.

Nick-seq thus provides an efficient, label-free approach to quantive mapping of DNA damage and modifications in genomes, with applications in DNA damage and repair, epigenetics, restriction-modification systems, and DNA metabolism.

**Methods – see the online Methods Section**

## Acknowledgements

The authors thank the MIT BioMicro Center, MIT Center for Environmental Health Science, Singapore-MIT Alliance for Research and Technology (SMART) for use of their facilities, and funding support from the National Natural Science Foundation of China (31630002), National Science Foundation of the USA (CHE-1709364), National Research Foundation of Singapore through the SMART Infectious Disease and Antimicrobial IRGs, National Institute of Environmental Health Sciences (P30-ES002109), and Fundamental Research Funds for the Central Universities of China (2015306020202). X.W. was supported by a fellowship from the China Scholarship Council (201606270163).

## Author Contributions

P.C.D. and B.C. designed Nick-seq. B.C., X.W., H.W. performed Nick-seq experiments. P.C.D, B.C., X.W., and J.Z. constructed the bioinformatics pipeline. M.S.D., C.G., D.Y., and L.W., contributed vital reagents. All authors discussed the results and contributed to the final manuscript.

## Data Availability

Sequencing data has been deposited in NCBI GEO database under accession numbers GSE138070, GSE138173, and GSE138476.

## Software and Code Availability

Custom scripts for processing the sequencing data are described in Methods and are available at https://github.com/BoCao2019/Nick-seq:. gitignore, NT_negative_strand.R, NT_positive_strand.R, TdT_negative+NT_positive.R, TdT_negative_strand.R, TdT_positive+NT_negative.R, and TdT_positive_strand.R.

## Supplementary Information

**Supplementary Figure 1.** Mapping Nb.BsrDI induced DNA strand-break sites in *E.coli* genomic DNA by Nick-seq. (a) The Nb.BsrDI cleavage motif derived from Nick-seq data. (**b**) A Venn diagram depicting the overlap of Nick-seq detected sites and Nb.BsrDI motif sites in the *E.coli* genome.

**Supplementary Figure 2.** Flanking sequence frequency analysis of H_2_O_2_-induced EndoIV-specific DNA damage sites based on Nick-seq data for genomic DNA (**a**) and plasmid (**b**) from cells exposed to a sublethal H_2_O_2_ dose. Site 0 represents a Nick-seq detected site.

**Supplementary Figure 3.** Detection of H_2_O_2_-induced EndoIV-specific DNA damage sites on genomic DNA (a) and an endogenous plasmid (b) in *E. coli* by Nick-seq.

**Supplementary Figure 4.** Distributions of H_2_O_2_-induced EndoIV-specific DNA damage sites on *E.coli* genomic DNA and plasmid. Outward from the center, circles represent: 0 and 0.2 mM H_2_O_2_ induced EndoIV-specific DNA damage sites.

**Supplementary Table 1**: Statistical analysis of paired-end DNA sequencing reads from Nick-seq.

**Supplementary Table 2**: Nb. BsmI sites identified by Nick-seq on the *E.coli* genome.

**Supplementary Table 3**: Nb. BsrDI sites identified by Nick-seq on the *E.coli* genome.

**Supplementary Table 4**: Phosphorothioate sites identified by Nick-seq on Salmonella genome.

**Supplementary Table 5**: AP sites identified by Nick-seq on *E.coli* genome.

**Supplementary Table 6**: AP sites identified by Nick-seq on plasmid in *E.coli*.

**Supplementary Table 7**: Regional distribution of AP sites identified by Nick-seq.

## Online Methods Section

### Materials

Nicking endonuclease Nb. BsmI, Nb. BspQI, Nb. BsrDI, Endonuclease IV, DNA polymerase I, OneTaq DNA polymerase, dNTPs, Nci I, Exonuclease III and RecJ_f_ were purchased from New England Biolabs. All DNA oligos were synthesized by Integrated Device Technology, Inc. (IDT). ddNTPs and alpha-thio-dNTPs were purchased from TriLink BioTech. Agilent Bioanalyzer 2100 was used for size analysis of DNA fragments. Other chemicals were of molecular biology grade. All cell lines used in this work are readily available from the authors.

### Cell growth and preparation of DNA

The PT-containing strain *Salmonella enterica* serovar Cerro 87 and its genomic DNA were prepared as described previously^9^. *E.coli* DH10B was used for nicking enzyme and H_2_O_2_-induced DNA damage mapping studies. A single colony of *E. coli* DH10B was grown in 5 mL LB medium overnight at 37 °C. 1mL cells were harvested by centrifuge at ambient temperature (unless indicated otherwise). Cells were resuspended and diluted with fresh 10 LB medium to a starting optical density at 600 nm (OD600) of 0.1, followed by growth at 37°C, 230 rpm until OD600=0.8 for DNA extraction or H_2_O_2_ treatment. 10 μL diluted H_2_O_2_ solution were added to the culture with a final concentration 0.1, 0.5, 1 and 2 mM. As un-exposed control, 10 μL sterile water was used instead of H_2_O_2_. After sitting at ambient temperature for 30 min, 10 μL of the cells were used for lethal dose (LD) analysis by counting the colony formation unit on LB agar plate. All the rest cells were harvested for DNA extraction with OMEGA bacterial genomic DNA or plasmid isolation kit by following the manufacture’s protocol.

### Mapping of modification/damage sites on DNA by NT-dependent method

These studies were initiated by random fragmentation of purified genomic DNA (1 μg) in each of three separate digestions with NciI, or HindIII and XhoI, or SalI, XbaI and NdeI. RNase A was also added to each reaction to remove trace of contaminating RNA. After digestion, the DNA was purified using a Qiagen PCR Purification Kit. The three purified DNA samples were mixed for the blocking step. Blocking of pre-existing strand-break sites was achieved in a reaction mixture (40 μl) containing 4 μl of reaction buffer (NEBcutsmart buffer), 1 μl of shrimp alkaline phosphatase (NEB), and 1 μg of template genomic DNA, with incubation at 37 °C for 30 min to remove phosphate at 3’ end of the strand-breaks. The phosphatase was then inactivated by heating at 70°C for 10 min. After cooling, 2 μl of ddNTPs (2.5 mM each, TriLink) and 1 μl of DNA polymerase I (10 U, NEB) was added to the reaction with incubation at 37 °C for 40 min to block any pre-existing strand-break sites. Shrimp alkaline phosphatase (1 μl) was then added at 37 °C for 30 min to degraded excess ddNTPs and the reaction was terminated by heating at 75 °C for 10 min. Following de-salting using a DyeEx column (QIAGEN), the DNA was ready for one of the following nick creation or conversion procedures.

Nicking *E. coli* DH10B genomic DNA with Nb. BsmI and Nb. BsrDI was accomplished in a 50 μl reaction mixture containing 1 μl Nb. BsmI (10 U, NEB), 1 μl Nb. BsrDI (10 U, NEB), 1 μg genomic DNA, and 1X NEBcutsmart buffer incubated at 65 °C for 1 h. The reaction was terminated by heating at 80 °C for 20 min and cooled down to 4 °C at the rate of 0.1 °C/s. The reaction product was used for NT- or TdT-reactions as described below with no further purification.

For mapping PT modifications, 40 μl of blocked DNA from *S. enterica* was mixed with 5 μl of dibasic sodium phosphate buffer (500 mM, pH 9.0) and 2 μl of iodine solution (0.1 N, FLUKA). After incubation at 65 °C for 5 min and cooling to 4 °C, the reaction product was purified using a DyeEX column (QIAGEN) to remove salts and iodine. The purified product was treated with shrimp alkaline phosphatase by adding 5 μl of NEBcutsmart buffer and 1 μl of phosphatase to remove 3’-phosphates arising from iodine cleavage. After incubation at 37 °C for 20 min and 75 °C for another 10 min, the product was kept on ice for the following NT- or TdT-reactions with no additional purification.

For mapping H_2_O_2_-induced AP sites, genomic DNA was extracted from H_2_O_2_-treated *E. coli* and the AP sites converted to strand-breaks in a 50 μl reaction mixture containing 1 μg genomic DNA, endonuclease IV (20 U), and 1X NEBcutsmart buffer, with incubation at 37 °C for 60 min. The reaction mixture was then kept on ice and used for the following NT- or TdT-reactions with no further inactivation or purification.

Following splitting of the nicked sample into two portions for NT- and TdT-reactions, the NT-reaction was achieved by further splitting the DNA sample into two parts: one for NT-reaction and the other as a negative control. NT-reaction was performed in a 50 μl reaction system containing 2 μl of α-thio-dNTPs (2.5 mM each, TRILINK), 1X of NEBcutsmart buffer, 2 μl of DNA polymerase I, and the DNA template. The negative control consisted of H_2_O instead of DNA polymerase I. The reaction mixture was incubated at 15 °C for 90 min and then terminated by heating at 75 °C for 20 min. The product was ready of template DNA digestion after the purification by DyeEx. The template DNA digestion reaction was performed in a 50 μl reaction system containing 200 U of exonuclease III, 5 μl NEBcutsmart buffer and DNA sample by incubating at 37 °C for 60 min. The DNA was then denatured by heating at 95 °C for 3 min and crashing on ice. RecJ_f_ (60 U) was then added to the reaction mixture with incubation at 37 °C for 60 min. For some DNA samples with high complexity of structure and/or modifications, digestion with an additional 60 units of RecJ_f_ might be necessary. After digestion, the enzymes were inactivated by incubation at 80 °C for 10 min. The DNA product was then purified using a Zymo Oligo Clean & Concentrator kit (Zymo) following the manufacturer’s protocol. The purified product was ready for Illumina library preparation.

Illumina library preparation was performed by the Clontech SMART ChIp-seq kit (Clontech) by following the manufacturer’s protocol. 12 cycles were used in the final step of PCR amplification. The PCR product of each sample was combined with its corresponding negative control and then size selected using AMPure XP beads (NEB). The purified library was submitted to Illumina NextSeq 500 instrument for 75 bp paired-end sequencing.

### Mapping of modification and damage sites by TdT-dependent method

The steps of DNA fragmentation, blocking, and nick conversion are the same as described above in NT-method.100 ng of the nick-converted DNA was denatured by heating at 95 °C for 3 min in 20 μl of H_2_O, followed by adding A poly(T) tail to the ssDNA in a 30 μl reaction system containing 3 μl DNA SMART buffer (Clontech), 1 μl Terminal Deoxynucleotidyl Transferase (TdT, Clontech) and 1 μl DNA SMART T-Tailing Mix (Clontech) by incubating at 37 °C for 20 min and terminating the reaction at 70 °C for 10 min. The primer annealing and template switching reaction was then performed with the Clontech SMART ChIp-seq kit (Clontech) by following the manufacturer’s protocol. The final step of PCR was perfomed using the Illumina primers provided in ChIp-seq kit and 12 cycles were used for amplification. The PCR product of each sample, with unique sequencing barcode, was combined with its corresponding negative control and then size selected using AMPure XP beads (NEB). The purified library was submitted to Illumina NextSeq 500 instrument for 75 bp paired-end sequencing.

### Data analysis

Sequencing results were processed on the Galaxy web platform (https://usegalaxy.org/). Initially, the paired-end reads were pre-processed by Trim Galore! to remove adapters, as well as trimming the first 3 bp on the 5’ end of read 1. All the reads were aligned to the corresponding genome using Bowtie 2. A custom method for peak calling of sequencing data was developed with BamTools, BEDTools and Rstudio. Briefly, the BamTools results were filtered based on R1 (selected for NT data) or R2 (selected for TdT data). The 5’ coverage (experiment sample and controls) or full coverage (controls) on each position were calculated based on the filtered BamTools results by BEDTools (positive and negative strand separately). For both NT or TdT data on each strand, three “.tabular” files containing the genome position and their corresponding read coverage (sample_coverage_5.tabular, control_coverage_5.tabular, control_coverage_full.tabular) were analyzed and exported from Galaxy after the previous BEDTools. These data were used to normalize the read coverage by the sequencing depth and then calculate the read coverage ratio of specific position compared to its up-downstream position in the same sample and the same sample in negative controls. Three ratios were calculated at each position by RStudio for modification site calling: coverage of position N (sample)/coverage of position N-1(sample), coverage of position N(sample)/coverage of position N+1(sample), and coverage of position N(sample)/coverage of position N(control). Positions with a ratio>1 were retained using the following R scripts: TdT_positive_strand.R TdT_negative_strand.R NT_positive_strand.R NT_negative_strand.R From these datasets, the intersection of the datasets from the NT and TdT methods were calculated using the following R scripts: TdT_positive+NT_negative.R TdT_negative+NT_positive.R The output files (CSV files; Excel format) contain the read coverage ratio information for the putative nick sites. The ratio cutoffs can be varied in the Excel spreadsheet as needed. For example, for site-specific nicking by Nb. BsmI and Nb. BsrDI, we determined that a ratio >2 was adequate to capture nearly all sites, while for variable sites (PT) or unknown samples (H_2_O_2_), the ratio was increased to 5-10.

