## Supplementary figures and images for "Nick-seq for single-nucleotide resolution genomic maps of DNA modifications and damage"

### Supplementary Figures

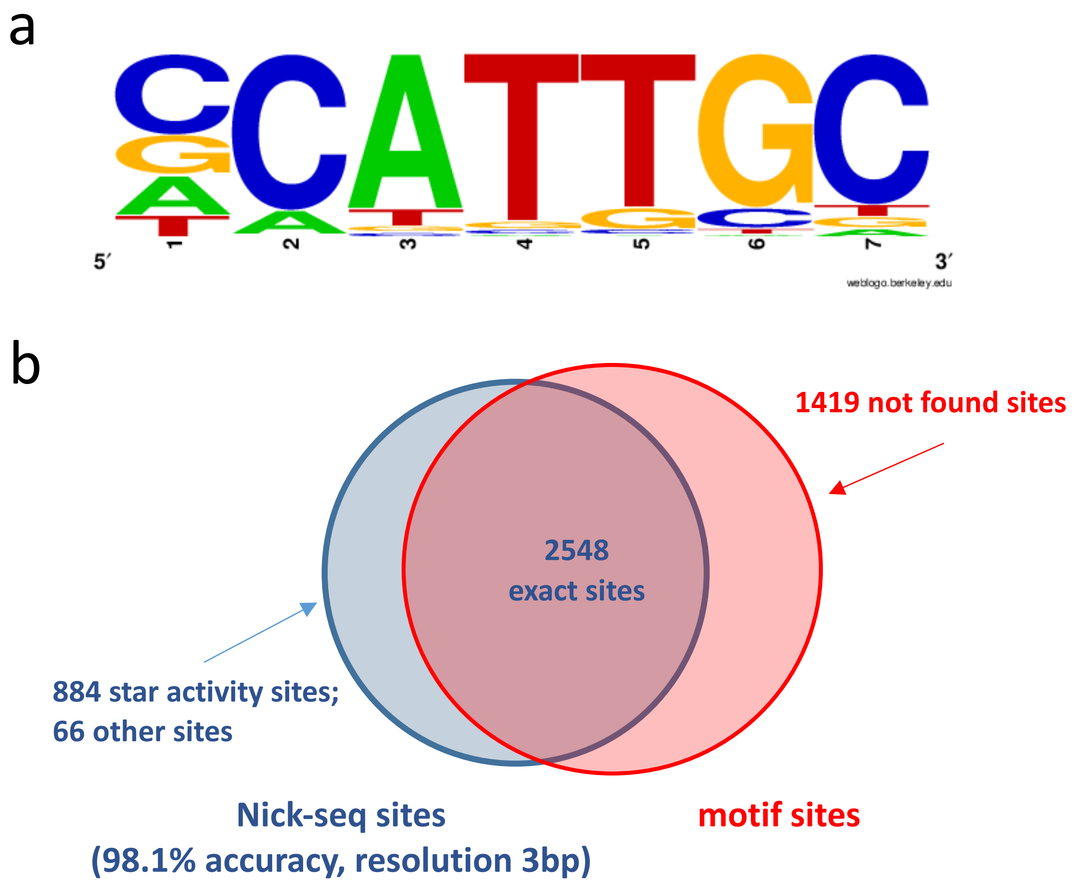

a

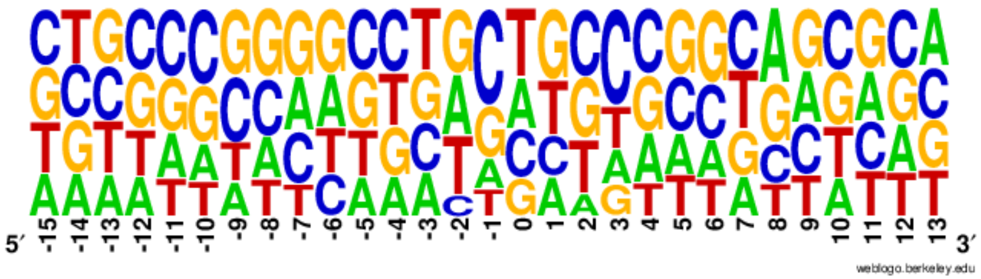


0.2mM $H_2O_2$-EndoIV

b

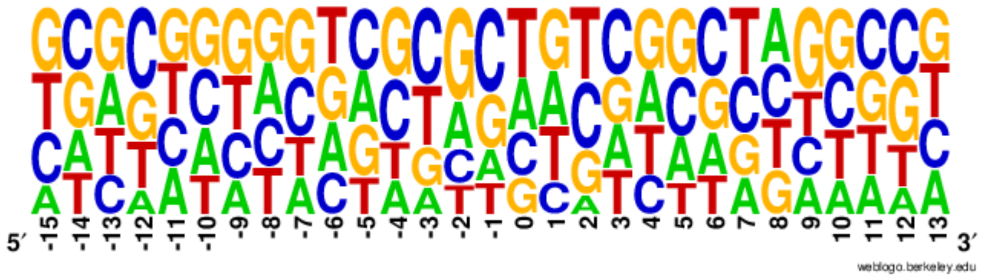


0.2mM $H_2O_2$-EndoIV-plasmid

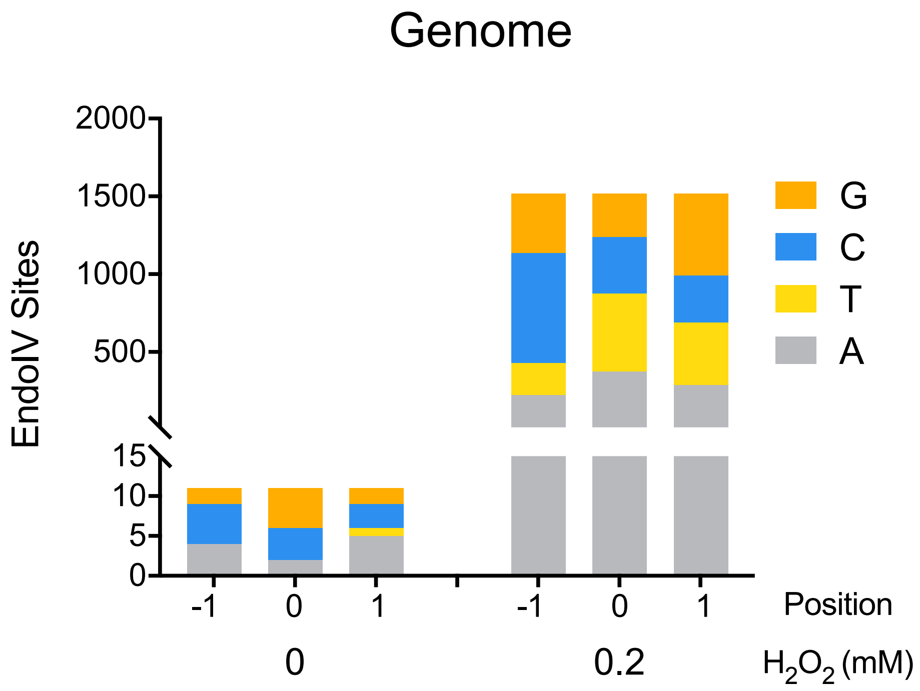

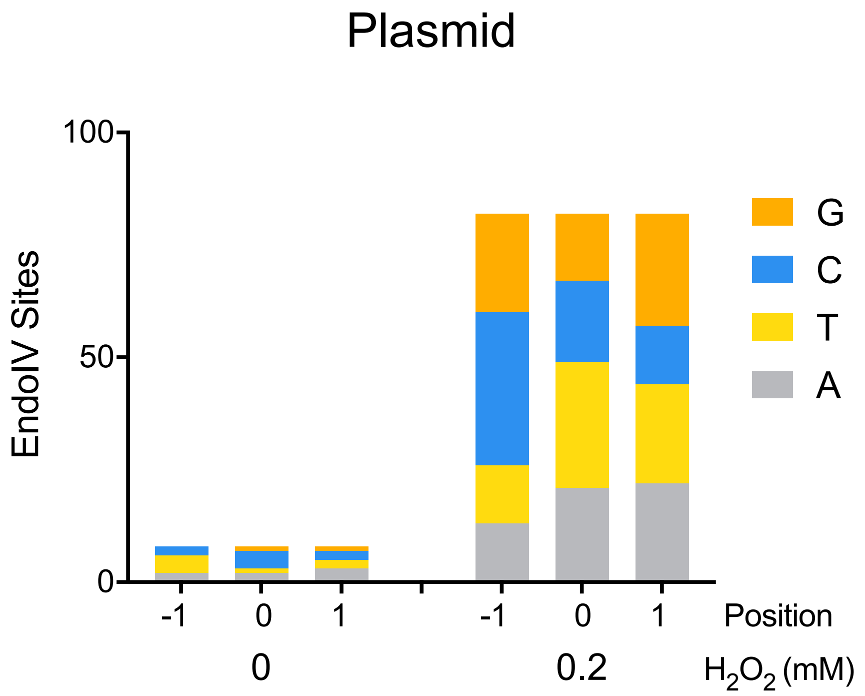

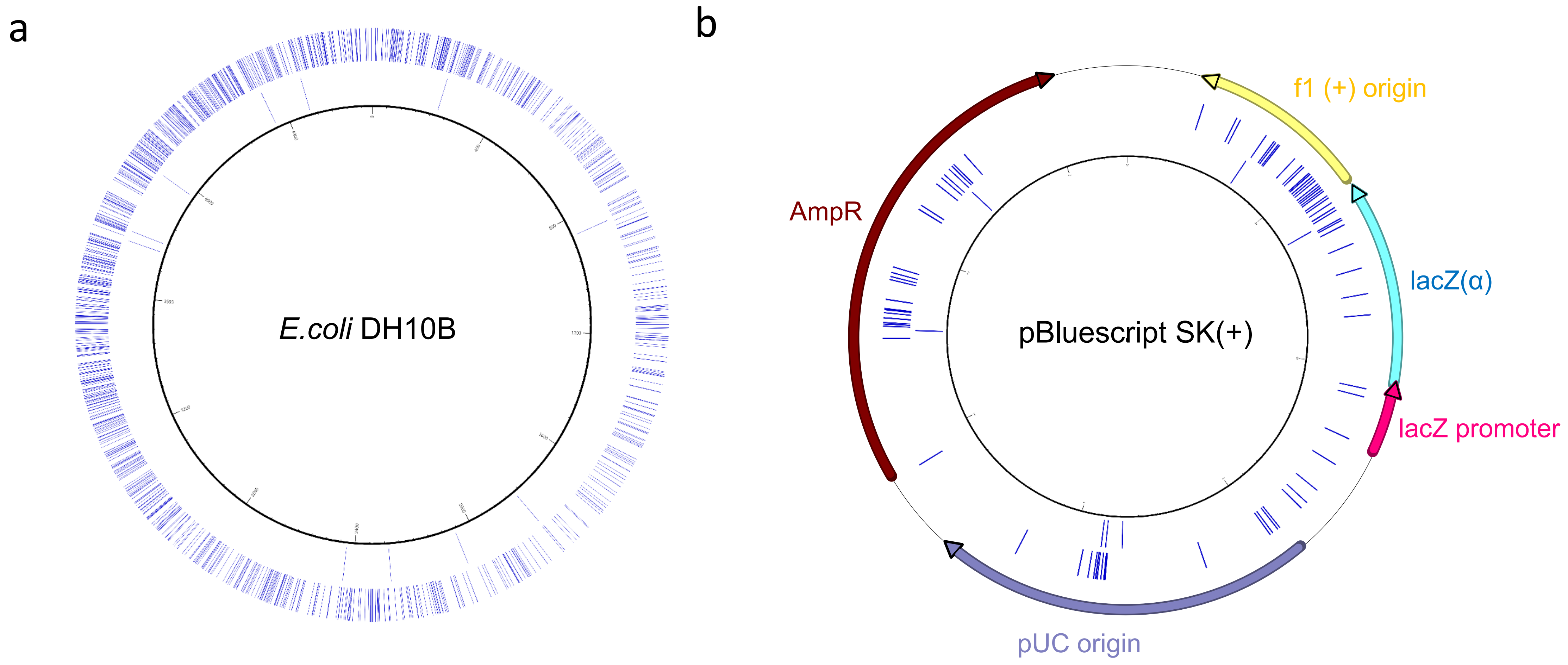
